# A transcription factor toggle switch determines differentiated epidermal cell identities in *Hydra*

**DOI:** 10.1101/2024.12.10.627691

**Authors:** Jaroslav Ferenc, Marylène Bonvin, Panagiotis Papasaikas, Jacqueline Ferralli, Clara Nuninger, Charisios D. Tsiairis

## Abstract

In *Hydra*, a simple cnidarian model, epithelio-muscular cells play a crucial role in shaping and maintaining the body architecture. These cells are continuously renewed as undifferentiated cells from the body’s mid-region get displaced toward the extremities, replacing shed, differentiated cells and adopting specific identities. This ongoing differentiation, coupled with the maintenance of distinct anatomical regions, provides an ideal system to explore the relationship between cell type specification and axial patterning. However, the molecular mechanisms governing epithelial cell identity in *Hydra* remain largely unknown. In this study, we describe a double-negative feedback loop between the transcription factors Zic4 and Gata3 that functions as a toggle switch to control epidermal cell fate. Zic4 is activated by Wnt signaling from the mouth organizer and triggers battery cell specification in tentacles. In contrast, Gata3 promotes basal disk cell identity at the aboral end. Functional analyses demonstrate that Zic4 and Gata3 are mutually antagonistic; suppression of one leads to the dominance of the other, and *vice versa*, resulting in ectopic cell specification. Notably, simultaneous knockdown of both factors rescues the phenotype, indicating that it is the balance between these transcription factors, rather than their absolute levels, that dictates cell identity. This study highlights the mechanisms by which distinct cellular identities are established at *Hydra* body termini and reveals how cell fate decisions are coordinated with axial patterning.

## Introduction

One of the key advantages of multicellularity is the division of labor among distinct cell types (Brunet and King 2017). Rather than housing conflicting functions within a single cell, multicellular organisms spatially segregate these functions into different regions of the body. This modularity has enabled the evolution of diverse body plans adapted to a wide range of environments, as seen in extant organisms (Valentine et al. 1994; Arendt et al. 2016). Consequently, understanding how cell fates are determined and spatially organized within the body is a central question in developmental biology.

Cell fate specification typically arises from a complex interplay between self-organization and pre-programmed events (Misailidis et al. 2021). Organizing centers, which emerge at multiple scales, from tissues to entire organisms, are key players in this process. Organizers direct the behavior of surrounding cells through chemical signals, often simultaneously influencing both cell identity and tissue morphogenesis (Anderson and Stern 2016). Interestingly, molecules associated with organizing centers are often evolutionarily conserved. A notable example is the Wnt signaling pathway, which establishes primary body axis polarity across most animals (Petersen and Reddien 2009; Loh et al. 2016).

*Hydra*, with its simple body plan and continuous cell differentiation, is an ideal model for studying the relationship between cell fate specification and spatial patterning (Vogg et al. 2019). This freshwater cnidarian has a tubular body organized along a single oral-aboral axis. The oral end consists of a mouth surrounded by tentacles for prey capture, while the aboral end terminates in a basal disk used for substrate attachment. Wnt signaling ligands are expressed around the mouth region (Hobmayer et al. 2000), and this region functions as an organizer. When transplanted into another individual, this tissue can induce the formation of a secondary body axis, composed primarily of host cells (Browne 1909). Such experiments are feasible because most *Hydra* cells are not terminally differentiated and remain responsive to organizer signals (Bosch et al. 2010).

The body wall of *Hydra* consists of two layers of multipotent epithelio-muscular cells: the epidermis and the gastrodermis. A third lineage, the interstitial cells (i-cells), resides between these layers. I-cells are fast-dividing and give rise to non-epithelial cell types such as neurons, gland cells, and nematocytes (stinging cells) characteristic of cnidarians (David 2012). Although both the epidermal and gastrodermal cells divide more slowly than i-cells, they do proliferate, particularly in the mid-body region (David and Campbell 1972). This ongoing division passively displaces cells that progressively move toward the extremities, where they undergo terminal differentiation (Dübel et al. 1987; Buzgariu et al. 2014) (Fig. 1A).

**Figure 1.**
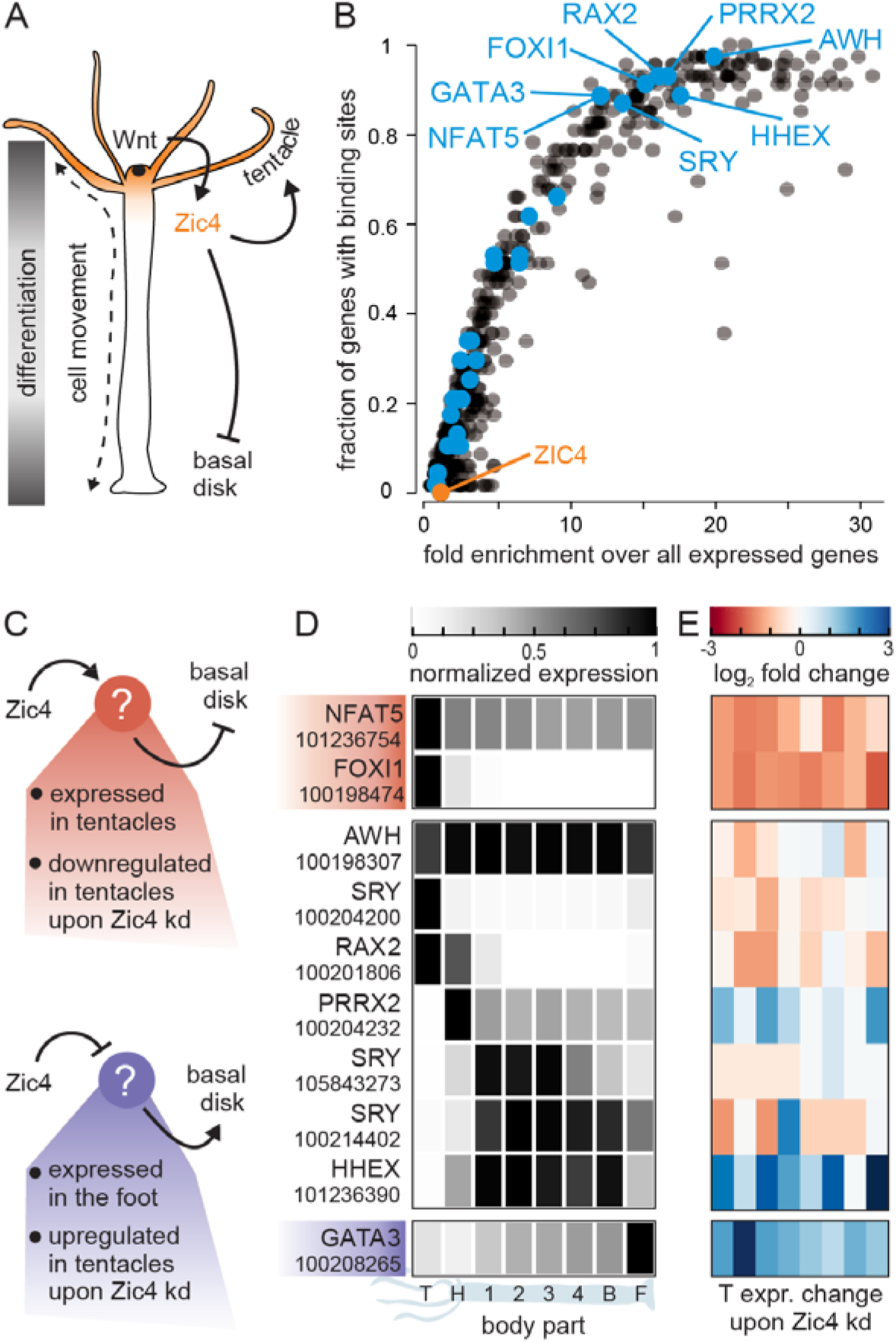
Potential downstream targets of Zic4 regulating basal disk identity. **(A)** Schematic of a *Hydra* body indicating expression domains of canonical Wnt ligands (black dot) and Zic4 (orange), and their effects on cell fate. For details see also Fig. S1. **(B)** Analysis of TF binding site enrichment in the promoters of foot-specific genes upregulated upon Zic4 KD. Each dot represents a single transcription factor. Transcription factors, whose promoters contain at least one *bona fide* Zic4 binding site, are highlighted in blue. **(C)** Possible modes of action for indirect inhibition of basal disk identity by Zic4. **(D)** Expression patterns of candidate intermediary factors from (B) along the body axis. Data from (28). **(E)** Expression changes in tentacles upon the downregulation of Zic4. Data from (20).

Differentiated epidermal cells exhibit striking morphological differences (Fig. S1). At the aboral end, basal disk cells have an inverted pyramidal shape, with a basally positioned nucleus and an abundance of peroxidase-containing granules near the apical surface (Davis 1973). At the oral end, epidermal cells, which pass around the organizer into the tentacles, differentiate into tentacle battery cells — a highly specialized cell type that attracts and houses several nematocytes in a complex with a sensory neuron (Hobmayer et al. 1990). These two structurally and functionally different cell fates are the endpoints of epidermal cell differentiation, and most epidermal cells eventually acquire one of them. However, how these alternative trajectories are regulated in connection with the axial extremities has long remained unclear.

Several genes have been identified as specifically expressed in or near the tentacles and basal disk, suggesting their involvement in specifying these structures. For example, *Wnt5* and *Wnt8* have been associated with tentacle development, while BMP ligands were linked to basal disk formation (Philipp et al. 2009; Wenger et al. 2019). However, the regulatory logic governing the choice between these two epidermal fates had been elusive. A key insight was provided by the recent identification of Zic4 as a crucial regulator of battery cell fate (Vogg et al. 2022). *Zic4*, a direct target of Wnt signaling, is expressed in the tentacles and at the tentacle base. Knocking down *Zic4* leads to an impressive phenotype: the transdifferentiation of tentacle battery cells into basal disk cells, suggesting that Zic4 plays a pivotal role in this fate decision. Nonetheless, the mechanism by which it prevents tentacle cells from adopting basal disk identity remains unclear. Does Zic4 directly repress basal disk-specific genes, or does it act through an intermediary factor?

In the present study, we addressed these questions by analyzing gene expression and *in silico* binding site predictions. We identified the transcription factor Gata3, previously implicated in basal disk formation, as the second component of a bistable toggle switch governing epidermal cell identities. Zic4 and Gata3 exhibit mutually exclusive expression patterns, and knockdown (KD) of *Gata3* results in the opposite phenotype of *Zic4* KD: the specification of battery cells at the aboral end. Remarkably, simultaneous KD of both transcription factors rescues the phenotypes, suggesting that cell fate is determined by the balance of Zic4 and Gata3, rather than their absolute expression levels. Our findings reveal a core regulatory switch that controls epidermal cell fate in *Hydra* and links these decisions to axial patterning.

## Results

### Identification of Zic4 downstream targets

Previous research has shown that the transcription factor (TF) Zic4 promotes tentacle fate in *Hydra* downstream of the canonical Wnt signaling from the head organizer (Vogg et al. 2022). Upon *Zic4* KD, battery cells begin to upregulate dozens of foot-specific genes, lose nematocytes, and ultimately transdifferentiate into basal disk cells. However, the mechanism by which Zic4 prevents the battery cell program from misexpression in normal tentacles remained unclear. To investigate this, we conducted a TF binding site enrichment analysis in the promoters of foot-specific genes that are overexpressed in tentacles following *Zic4* KD (Fig. 1B and Supplementary file 1). Surprisingly, Zic4 binding sites showed no enrichment, implying that another intermediary factor might be involved. Potential candidates for this function could be found among TFs with highly enriched binding sites in the dataset, which are themselves Zic4 targets. We therefore examined the promoters of genes encoding these TFs to identify *bona fide* Zic4 binding sites, which led us to pinpoint eight TFs (Nfat5, Gata3, Sry, FoxI1, Hhex, Rax2, Prrx2, and Awh) as probable Zic4 targets that could also regulate most foot-specific genes (Fig. 1B, blue dots).

A Zic4-dependent effector could maintain battery cell identity in two ways: it might be upregulated by Zic4 and inhibit the basal disk program, or it might be downregulated by Zic4 and activate this program (Fig. 1C). In either case, Zic4 would ultimately prevent basal disk cell differentiation, and its absence would lead to ectopic foot identity establishment in tentacles. Using RNA-seq data of positional expression profiles (Ferenc et al. 2021), and changes in tentacles upon *Zic4* KD(Vogg et al. 2022), we narrowed down our shortlist to three candidate regulators of basal disk identity downstream of Zic4: Nfat5, FoxI1, and Gata3 (Fig. 1D-E). Nfat5 and FoxI1 are mainly expressed in tentacles and downregulated upon *Zic4* KD. *Gata3* expression gradually increases along the body axis, peaking in the foot. Upon *Zic4* KD, *Gata3* expression is upregulated in tentacles. Based on whole-animal scRNA-seq data (Siebert et al. 2019), *Nfat5* expression does not appear to be strongly correlated to Zic4 levels in individual cells. However, *FoxI1* expression shows a strong positive correlation with *Zic4*, while *Gata3* levels are strongly anticorrelated, suggesting regulatory relationships (Fig. 2A, S2 and Table S1). FoxI1 could thus act as a negative regulator of basal disk fate, while Gata3 may function as a positive one. While FoxI1 may help repress foot-specific genes in tentacle cells, it is unlikely that fate acquisition relies only on the absence of this factor. Consequently, we focused on Gata3, the candidate positive regulator, which was also previously shown to be essential for foot regeneration (Ferenc et al. 2021).

**Figure 2.**
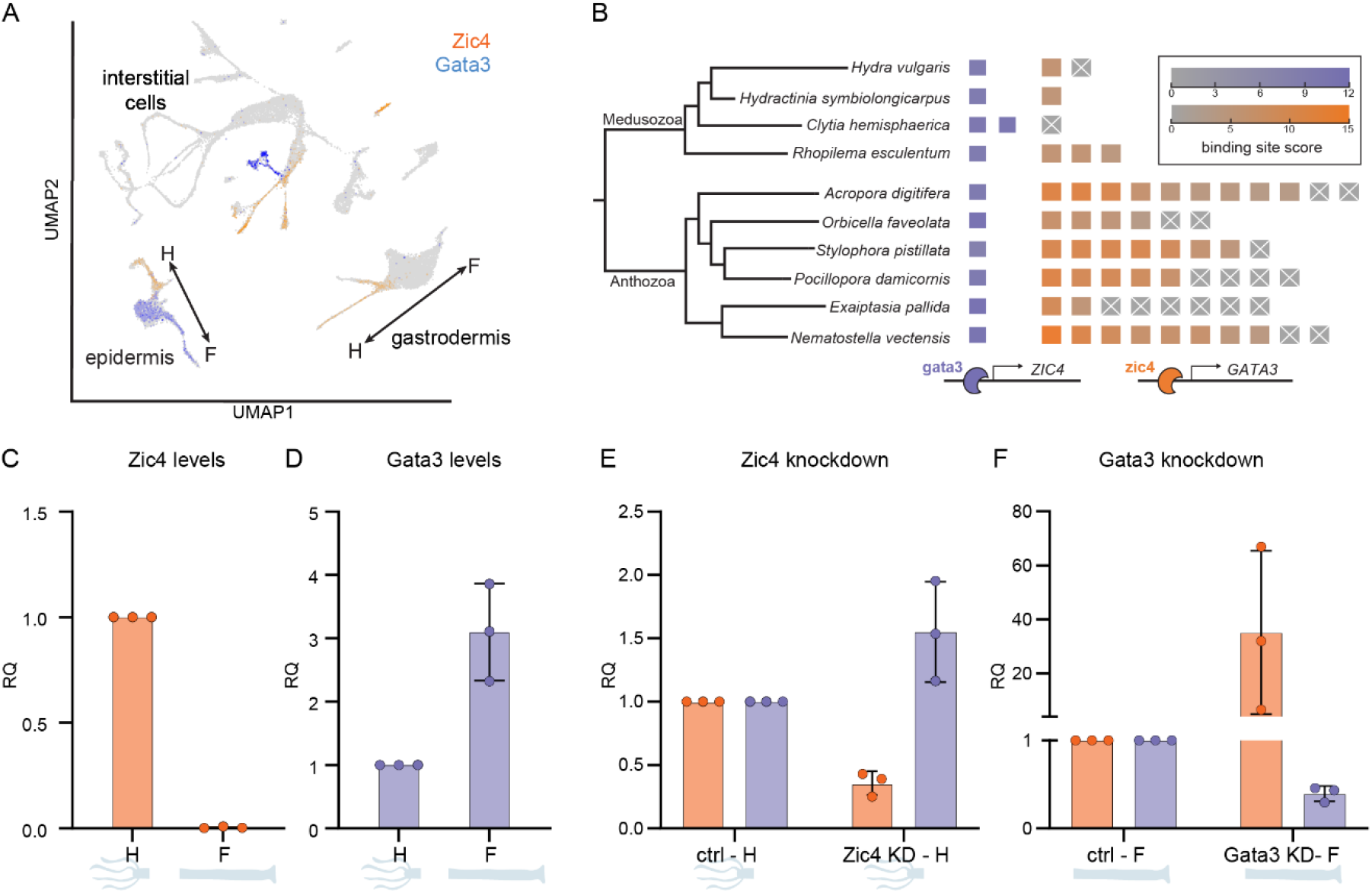
Mutually exclusive expression of Zic4 and Gata3. **(A)** Expression profiles of Zic4 (orange) and GATA (blue) in the *Hydra* single cell atlas (research.nhgri.nih.gov/HydraAEP). Note the diverging trajectories in ectoderm and the i-cell lineage. Arrows indicate the direction of the axis between the two poles: oral (head = H) - aboral (foot = F). **(B)** Binding site conservation across cnidarians. Gata3 binding sites on *Zic4* promoters in purple, and Zic4 binding sites on *Gata3* promoters in orange. Each square represents a transcript isoform(s) with a unique transcription start site (TSS). Color intensity indicates the motif score for the strongest binding site detected 1.5kb upstream of each TSS. Tree topology based on NCBI taxonomy **(C-F)** Quantitative PCR analysis of *Zic4* (orange) and/or *Gata3* (purple) transcript levels in the oral (H) and aboral (F) ends of the body axis. **(C-D)** Homeostatic animals. **(E-F)** *Zic4* or *Gata3* RNAi. In (C) to (F), each data point represents one biological replicate and error bars indicate standard deviations. ctrl - control, KD - knockdown, H - head, F - foot.

### Mutual regulation of Zic4 and Gata3

The strong anticorrelation between *Zic4* and *Gata3* expression in single cells (Fig. 2A, S2A-B,E) suggests a mutually exclusive regulatory relationship possibly important for specifying the respective cell types. This hypothesis is further supported by trajectories in the scRNA-seq data, where cells in the epidermal lineage exhibit a clear divergence in *Zic4* and *Gata3* expression. Cells differentiating into oral fate upregulate *Zic4*, while those differentiating into basal disk show higher *Gata3* transcript levels (Fig. 2A). Interestingly, a similar binary choice pattern appears in certain neuronal lineages as well. A recent study demonstrated the involvement of Gata3 in specifying neurons at the aboral end (Primack et al. 2023), suggesting that this cross-regulation between Zic4 and Gata3 may extend beyond the epidermal lineage. To explore whether this regulatory pattern might be conserved in other species, we analyzed binding sites in the promoters of *Zic4* and *Gata3* homologs across ten cnidarians (Fig. 2B and Supplementary file 2). At least one *bona fide* Gata3 binding site was found within 1.5 kb upstream of the transcription start site (TSS) in *Zic4* homologs across all species. Notably, *Gata3* homologs tend to have multiple transcript variants with differing TSS regions, and not all possess a strong Zic4 binding site, suggesting this diversity may allow for more nuanced regulation. However, except for *Clytia, Gata3* homologs in all other studied species contain at least one transcript isoform with a Zic4 binding site upstream of its TSS. This regulatory motif is thus likely functional across the entire cnidarian phylum.

To directly assess mutual inhibition between these factors in *Hydra*, we conducted KD experiments paired with quantitative PCR (q-PCR) on specific body parts. Consistent with previous RNA-seq data, *Gata3* is predominantly expressed in the foot and significantly upregulated in tentacles following *Zic4* KD (Fig. 1E). Q-PCR performed on dissected oral and aboral body regions confirmed this finding (Fig. 2C-D). We then tested the effect of RNAi-mediated downregulation of *Gata3* or *Zic4* on the expression levels of each gene at the axis ends. *Zic4* RNAi led to a notable decrease in *Zic4* expression in the head region, accompanied by an increase in *Gata3* levels (Fig. 2E), aligning with RNA-seq observations. Conversely, *Gata3* RNAi reduced *Gata3* transcripts and induced *Zic4* expression in the aboral region (Fig. 2F). Together, our data reveal that Zic4 and Gata3 form a double negative feedback loop that is likely conserved across diverse cnidarian species.

### Gata3 depletion leads to ectopic battery cell specification

The double-negative feedback loop between Zic4 and Gata3 suggests a toggle-switch mechanism where terminal cell identity is determined by the dominant factor. Thus, when Zic4 is depleted, Gata3 dominates in the tentacles, leading to ectopic basal disk specification. Conversely, depleting Gata3 should allow Zic4 to dominate in the foot region, causing ectopic battery cell specification. Given our previous findings that Gata3 is essential for foot regeneration (Ferenc et al. 2021), we sought to assess whether *Gata3* KD would cause cell identity shifts at the amputation site rather than simply inhibiting regeneration. To do so, we combined peroxidase activity detection, to mark the basal disk, with phalloidin staining to reveal cell morphology. In *Gata3* KD regenerates, basal disk peroxidase activity was absent, while epithelial cells appeared larger and contained multiple nematocytes, indicated by the characteristic actin staining of the cnidocil (Golz and Thurm 1991; González Agosti and Stidwill 1992) (Fig. S1). This phenotype is consistent with a shift to battery cell-like fate (Fig. 3A, Fig. S4).

**Figure 3.**
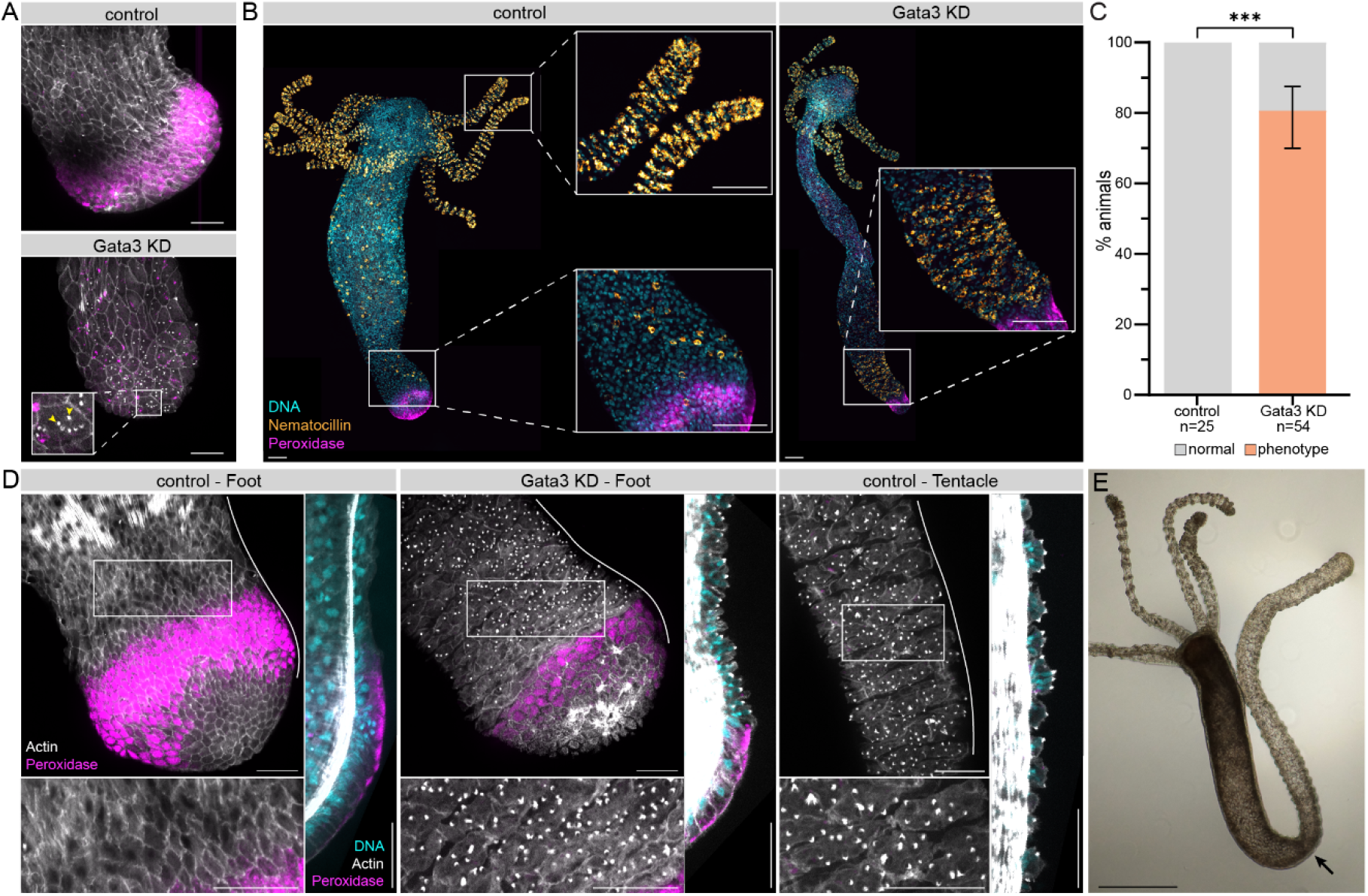
Ectopic battery cell specification upon *Gata3* knockdown. **(A)** Regenerating feet 3 days post amputation. Maximum intensity projections of confocal images of the epidermal layer. Peroxidase activity in magenta, actin in grey. Control (*GFP* RNAi), representative of 7/7 samples. *Gata3* RNAi, representative of 8/10 samples. Note the stained cnidocils (arrowheads). **(B)** Maximum projection confocal images of whole-mount *Hydra* after *GFP* RNAi (control) or *Gata3* RNAi. *Nematocilin* (orange), peroxidase activity (magenta) and DNA (cyan). **(C)** Quantification of animals with ectopic *Nematocilin* in the foot. Percentages are shown for n animals pooled from at least 2 independent replicates. Error bar indicates minimum and maximum values across individual replicates. Data was analysed using Fisher’s exact test. \*\*\**P*<0.001. **(D)** Maximum projection confocal images of actin (grey) and peroxidase staining (magenta) in the epidermins after *GFP* (control, representative of 5/5 foot and tentacle samples) or *Gata3* RNAi (representative of 5/7 samples). Rectangles indicate areas magnified below the projections. Lines indicate midline optical sections shown next to the projections. DNA channel (cyan) added to the optical sections. Note the accumulation of nematocytes above the foot and the morphological correspondence with a control tentacle. **(E)** Brightfield image of a *Gata3* RNAi animal showing a strong phenotype with a tentacle-like structure instead of a foot (arrow indicates the beginning of the ectopic tentacle). Representative of 41/105 samples. Scale bars (main panels and insets): A, D: 50 μm, B: 100 μm, E: 500 μm.

We then investigated whether a similar phenotype could emerge in a homeostatic setting, where animals are maintaining an existing foot rather than building one *de novo*. We used hybridization chain reaction RNA-fluorescent in situ hybridization (HCR RNA-FISH) with probes for *Nematocilin*, a nematocyte-specific transcript (Vogg et al. 2022). In control animals, *Nematocilin* signal is concentrated in the tentacles, with a salt-and-pepper pattern across the body, representing relatively scarce body wall nematocytes. However, in *Gata3* KD animals, we detected patches of densely concentrated nematocyte staining, reminiscent of tentacles, directly above the foot (Fig. 3B). This phenotype, absent in controls, was highly penetrant in *Gata3* KD animals, consistently appearing across replicates (Fig. 3C). Morphologically, the cells in these clusters were indistinguishable from battery cells (Fig. 3D). While the apical surface of cells adjacent to the foot is elongated parallel to the body axis, transformed cells in *Gata3* KD animals elongate perpendicularly, much like battery cells. These cells are also notably larger and contain numerous nematocytes, as shown by multiple nuclei and cnidocil staining. In some cases, the phenotype is remarkably strong, leading to the development of a long tentacle-like structure at the aboral end (Fig. 3E, Fig. S5A-B).

Interestingly, despite ectopic differentiation of battery cells above the existing basal disk, we never observed a complete loss of basal disk markers (such as peroxidase) upon *Gata3* KD even though the signal often appears reduced (Fig. 3D). This persistence of basal disk identity at the aboral end suggests that, unlike battery cells, basal disk cells can retain their differentiated state even at low levels of its crucial positive regulator Gata3. Incoming cells, however, respond to the altered Zic4/Gata3 balance as predicted, ectopically adopting a battery cell fate.

### Simultaneous downregulation of Gata3 and Zic4 rescues ectopic cell specification

As shown above, individual knockdowns of either *Zic4* or *Gata3* alone are sufficient to drive ectopic differentiation directed by the opposing TF (Fig. 4A-B). Since these two factors operate in a double-negative feedback loop, it should be possible to rescue both phenotypes by simultaneously downregulating both TFs. Given a comparable RNAi strength for both genes (Fig. S3), this approach would likely reestablish the normal balance between Zic4 and Gata3, similar to what is observed in control animals, albeit at lower absolute levels. Indeed, after performing the combined knockdown, we achieved a full rescue of the ectopic basal disk phenotype (Fig. 4A and C). In some animals, tentacles still appeared slightly shorter but were otherwise morphologically normal. Likewise, the ectopic battery cells that typically appear next to the foot following a single *Gata3* KD were absent in most double KD animals. Although small, transformed patches occasionally persisted, we never observed a strong phenotype involving widespread transformation of the body wall (Fig. 4B and D, Fig. S5B). These differences probably arose as a result of variable RNAi efficiency.

**Figure 4.**
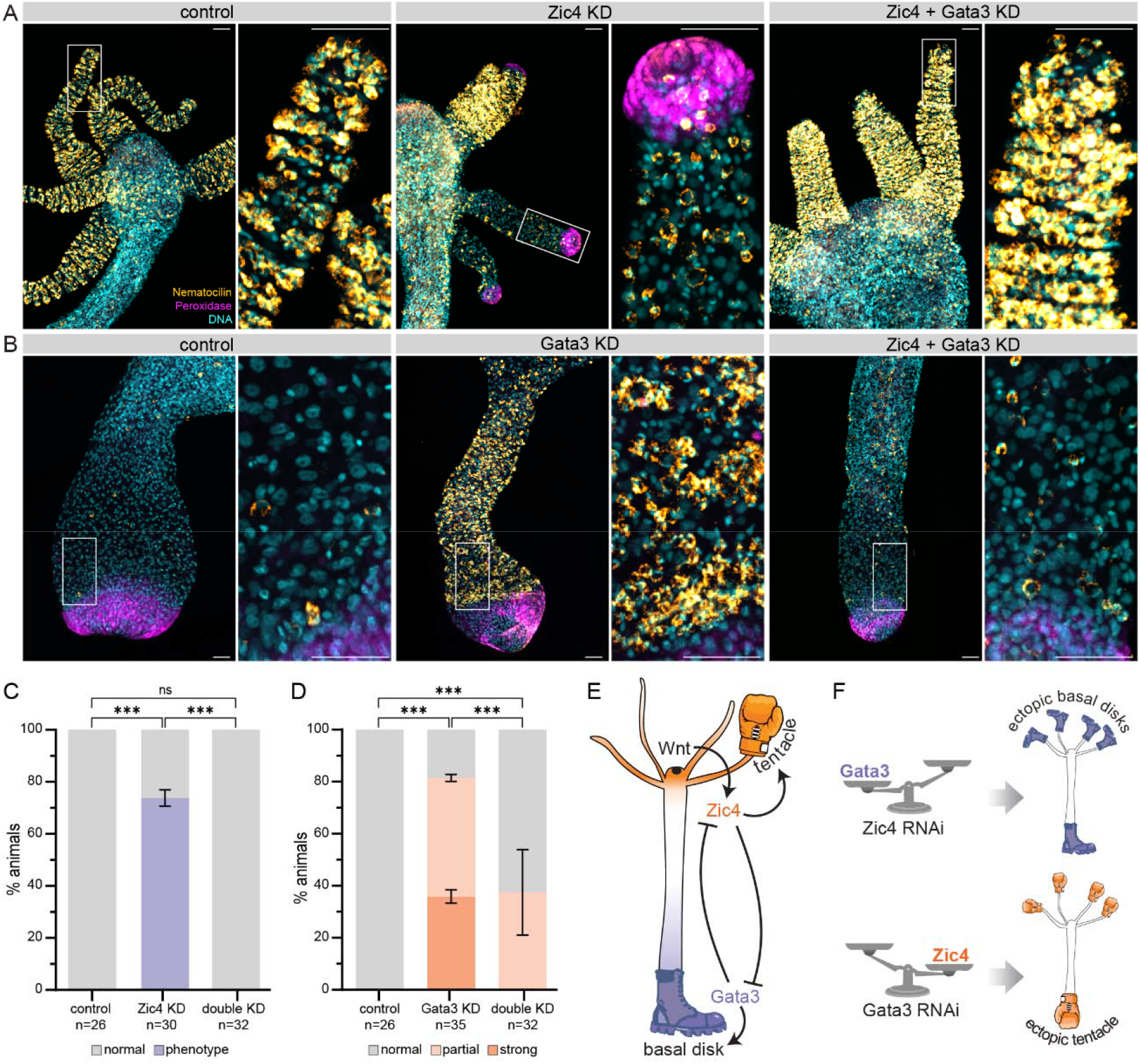
Phenotype rescue upon *Zic4* and *Gata3* double knockdown. **(A-B)** Maximum projection confocal images of oral (A) and aboral (B) ends of animals after *GFP* RNAi (control), *Zic4* or *Gata3* RNAi and *Zic4* + *Gata3* RNAi. DNA in cyan, peroxidase activity in magenta, and *Nematocilin* in orange. Panels on the right show closeups of areas highlighted by rectangles. Scale bars (main panels and closeups) 100μm. Note the ectopic peroxidase staining in tentacle tips upon *Zic*4 RNAi, and ectopic *Nematocilin* staining adjacent to the foot upon *Gata3* RNAi, as well as rescue of both phenotypes in the double KD. **(C)** Percentage of animals with ectopic peroxidase staining in tentacles upon *Zic4* KD and double KD. **(D)** Percentage of animals with ectopic *Nematocilin* staining adjacent to the foot in *Gata3* KD and double KD. For examples of strong and partial phenotypes refer to Fig. S5. Data in (G) and (H) shown for n animals pooled from at least 2 independent replicates. Data was analysed using Fisher’s exact test. \*\*\**P*<0.001. **(E-F)** Schematic of proposed model for controlling the choice between basal disk and battery cell fate in *Hydra*. At the oral pole, Wnt signaling (black dot) activates Zic4 (orange) expression and thus basal disk identity. Without it, Gata3 dominates at the opposite end, leading to basal disk fate establishment (E). Experimentally depleting one of the factors results in ectopic cell fate acquisition at the opposite body end (F).

Our findings reveal a double-negative feedback loop between Zic4 and Gata3, which is crucial for determining two distinct epidermal identities — battery cells in the tentacles and basal disk at the aboral end of a *Hydra* body. As cells migrate towards the body extremities, the expression of one factor gradually takes precedence, ultimately specifying the appropriate terminal cell identity (Fig. 4E). Consistent with the role of this regulatory motif as a toggle switch, experimentally manipulating the levels of either factor leads to predictable ectopic differentiation outcomes (Fig. 4F). However, restoring the balance through a simultaneous knockdown rescues both phenotypes, highlighting the importance of their ratios rather than absolute levels for cell fate decisions.

## Discussion

The cellular events involved in constructing and maintaining a simple *Hydra* body have been subject to intensive study for decades. Thanks to these investigations, the contributions of cell division, migration, differentiation, and death are now relatively well understood (Vogg et al. 2021). More recently, molecular studies have complemented this cellular perspective. In particular, the advent of single-cell sequencing has illuminated the transcriptional signatures of various cell types, which were previously defined largely by morphology, also offering hints of candidate regulators that specify them (Siebert et al. 2019; Cazet et al. 2023). However, mechanistic studies remain sparse, especially for the myoepithelial lineages.

Building on our previous work, which identified the transcription factor Zic4 as a key regulator of battery cell fate (Vogg et al. 2022), we have now discovered that Gata3 also plays a crucial role in controlling the terminal identities of epidermal cells in *Hydra*. Our data indicate that these two factors function together in a double-negative feedback loop, controlling the decision between basal disk and battery cell identities. KD of either factor allows the other to dominate, leading to a corresponding change in cell identity. However, simultaneous KD of both factors results in largely normally differentiating cells, suggesting that the balance between them, rather than absolute levels, is critical for the regulation. Thus, Zic4 and Gata3 represent terminal selectors (Arendt et al. 2016; Hobert 2016) that govern transcriptional programs and identities of differentiated *Hydra* epidermal cells, as shown by the cell identity changes resulting from manipulating their expression experimentally (Arlotta and Hobert 2015).

Interestingly, the downregulation of Gata3 only affects the identity of undifferentiated cells moving towards the foot region, without impacting existing basal disk cells. In contrast, Zic4 downregulation influences both precursor and differentiated battery cells, causing the latter to transdifferentiate into basal disk cells (Vogg et al. 2022). This suggests that the basal disk identity is more stable and/or represents a default fate for epidermal cells in *Hydra* in the absence of specific signals such as Wnt ligands from the mouth organizer. As shown previously, canonical Wnt signaling promotes *Zic4* expression and thereby the battery cell fate (Vogg et al. 2022). The organizer’s presence at the oral pole also partly explains the segregation of epidermal identities along the body axis. This situation is reminiscent of early zebrafish hematopoiesis, where a conserved transcription factor toggle switch between Pu.1 and Gata1 is also differentially regulated along the main embryo axis (Rhodes et al. 2005). Posterior signals promote *Gata1* expression, leading to erythroid fate, while anterior signals increase *Pu*.*1* expression, promoting myeloid fate, resulting in spatial segregation of progenitors. In *Hydra*, it remains to be determined whether there are specific foot-inducing signals (such as BMP signaling as proposed by some authors (Wenger et al. 2019)), which could stimulate *Gata3* expression, or if the effect results from an asymmetric switch (Lyons et al. 2014) defaulting to aboral identity the absence of oral factors. Nevertheless, since only the body extremities respond to altered Zic4/Gata3 ratios, cells likely perceive other cues in the terminal regions that are a prerequisite for differentiation.

To ensure proper development, changes in cell identity must not only occur in the correct body regions but also in coordination with morphogenetic events. The alterations in cellular properties resulting from identity shifts can, in turn, influence tissue morphogenesis (Chan et al. 2017; Collinet and Lecuit 2021). Interestingly, in animals exhibiting the strongest phenotype following *Gata3* KD, the foot portion of the body column becomes elongated and tentacle-like. We speculate that such morphology might result from the geometry of battery cells rather than active tissue remodeling in response to identity change. Similarly, the shortened tentacles observed upon *Zic4* KD, where existing battery cells transdifferentiate to a basal disk fate, could arise in the same way. Nevertheless, manipulating neither Zic4 nor Gata3 led to complete loss of tentacles or gain of ectopic ones, as is the case of upstream regulators (Hobmayer et al. 2000; Broun et al. 2005). This observation is consistent with their role as cell fate switches rather than morphogenesis regulators *per se*.

Since core regulatory complexes often exhibit combinatory logic when specifying different cell fates (Hobert 2008; Allan and Thor 2015), it is noteworthy that *Zic4* and *Gata3* are not exclusively expressed in epidermal cells. Single-cell RNA sequencing data reveal a similar bifurcating expression trajectory on the way to neuronal subpopulations, suggesting that this switch might additionally specify neuronal identities in combination with other factors (Primack et al. 2023). Identifying these factors will be an interesting avenue for future exploration. Binding site analysis of *Zic4* and *Gata3* promoters also points out that this switch is not unique to *Hydra* but shows wider conservation across cnidarians with the exception of *Clytia*. In this organism, we detected no bona fide Zic4 binding site in the *Gata3* promoter. Since, however, *Clytia* only appears to have one *Zic* gene in the genome, it is possible that while maintaining cross-regulation, the sequence that it binds is too divergent to be detected in our analysis. Interestingly, a recent study comparing cell type transcriptional profiles (Cazet et al. 2023) revealed that the basal disk and peduncle cells of *Hydra* are homologous to the exumbrella epidermis of *Clytia* medusa. No such correspondence was established for the battery cells, however, which might also indicate a relevant change in the underlying regulatory logic. Notably, in anthozoans, the *Gata3* homologs have a higher number of transcript variants compared to medusozoans. Many of these variants differ by the position of the transcription start site, not always containing a Zic4 binding site in the upstream region, which hints at further diversification of regulatory possibilities among these different variants. Moving forward, it will be interesting to verify and compare the function of the Zic4/Gata3 switch across species and gain further insights into cell type homology in cnidarians.

Taken together, our findings provide new insights into the mechanisms behind cell identity in *Hydra*. Thanks to its simple anatomy and evolutionary position, this animal has been a fruitful model system for understanding the principles of building bodies that are common to all metazoans (Vogg et al. 2019). By identifying and characterizing the toggle switch between Zic4 and Gata3, we elucidated the molecular logic governing the decision between two major epidermal cell fates and their spatial connection to body axis poles. Integrated with other datasets, and future work, our results will contribute to the understanding of gene regulatory networks governing cell identities and shaping the simple body of this model early-branching metazoan with possible implications going beyond cnidarian biology.

## Materials and Methods

### Animal strains and culture conditions

All experiments were performed using either the *Hydra vulgaris* 105 strain or derivatives of the AEP strain (AEP, AEP sexy, Ecto::GFP, Zic4::GFP). Animals were kept in *Hydra* medium (HM, 1 mM CaCl_2_, 0.2 mM NaHCO_3_, 0.02 mM KCl, 0.02 mM MgCl_2_, and 0.2 mM tris-HCl (pH 7.4)) at 18 °C and fed with freshly hatched *Artemia* nauplii three times a week. Individuals without buds, starved for at least 24 hours, were used for experiments if not indicated otherwise. All experiments were performed in accordance with the Swiss national regulations for the use of animals in research.

### Scoring foot regeneration in bisected animals

Animals were bisected at 50% body length, and the halves were kept separately in the wells of 6-well plates at 18°C. Fixation and peroxidase staining or *ISH* with foot marker crim-like, were performed 3 or 6 days after bisection on the head halves.

### Peroxidase foot staining

Fluorescent peroxidase staining was performed based on previously published method for chromogenic staining (Hoffmeister and Schaller 1985). Animals were relaxed in 2% urethane in HM and fixed with 4% paraformaldehyde (PFA) in HM at 4°C overnight. Animals were then washed in phosphate-buffered saline + 0.1% Tween 20 (PBST) for 15 min and stained for 30 min in the staining solution (1x Alexa Fluor488 tyramide reagent (#B40953, Invitrogen), 0.03% H_2_0_2_, 0.25% Triton-X diluted in PBS) for 30 min at RT in the dark with gentle agitation. To stop the enzymatic reaction, samples were washed once again in PBST for 15 min. To stain for actin and DNA, animals were incubated for 30 min in PBST containing Hoechst 33258 (1:1000, #94403, Sigma Aldrich) and Phalloidin Atto 565 (1:50, #94072, Sigma Aldrich). Animals were then washed several times with PBST and mounted for imaging using ProLong Gold Antifade mountant (#P36930, ThermoFischer).

### Hybridization chain reaction RNA-FISH

The HCR RNA-FISH protocol has been adapted for *Hydra* based on the manufacturer’s instructions (Molecular Instruments) and was performed according to previously published protocol (Vogg et al. 2022). For some experiments, this protocol was combined with peroxidase staining, in such cases minor modifications were made as described below. All incubations were performed at RT, unless stated otherwise. Animals were relaxed in 2% urethane in HM for 1 min and fixed for 1h in 4% PFA at RT. After fixation, the samples were washed 2x 10 min in PBST. If samples were to be stained for peroxidase activity, the staining was performed at this point as detailed above. Samples were rinsed once and washed once 5 min with PBST before continuing with the HCR protocol, performing all incubation and wash steps in the dark. Samples were incubated in 250μl of HCR probe hybridization buffer, without probe, for 30 min at 37°C. Samples were then incubated with 250 μl of HCR probe hybridization buffer containing 1 pmol of *Nematocilin* probe (4nM) overnight (16 hours) at 37°C in the dark. The probe was synthesized by Molecular Instruments (lot number RTC726, binds to both *NematocillinA* (100192251) and B (100192252)). Hybridization probes were washed out with 5 washes of 1h in 500μl of HCR probe wash buffer at 37°C, followed by two washes of 5min with a 5× sodium chloride sodium citrate/0.1% Tween 20 (5xSSCT) solution. Signal was pre-amplified with 250 μl of amplification buffer for 30min. 7.5 pmol of amplifying hairpin RNAs, provided by Molecular Instruments, were heated at 95°C for 1min 30s and allowed to cool down in the dark for 30 min at RT. Samples were incubated with 250μl of amplification buffer containing the two hairpins (0.03μM) overnight (16 hours) in the dark. Samples were washed in 5×SSCT solution in the dark: 2x 5 min, 2x 30 min, and 1x 30 min with 5×SSCT containing 4′,6-diamidino-2-phenylindole (DAPI, 1μg/ml, #62248, ThermoFischer) and/or Phalloidin Atto 565 (1:50, #94072, Sigma Aldrich), and 1x 5min in 5xSCCT. Samples were carefully mounted on a slide with ProLong Gold Antifade mountant (#P36930, ThermoFischer) for imaging.

### Image acquisition

Brightfield images were acquired with an hybrid microscope Revolve from Echo (RVL2-K), using the inverted mode with a 4.0x UPlanXApo objective and a transmission light camera (CMOS-colour).

Fluorescent imaging was conducted using spinning disk confocal microscopes. Whole-mount animals stained for Nematocillin probe, peroxidase activity and nuclei were imaged using an inverted motorized stand AxioObserver7 (Zeiss) equipped with CSU-W1 Confocal Scanner Unit (Yokogawa) with a 50 um pinhole disk unit, 2 black-illuminated sCMOS cameras (Prime 95B model, Photometrics) and controlled with VisiView 6.0 imaging software. Illumination was achieved with 3 excitation lasers operating at 405nm, 488nm and 640nm. A Plan Apochromat air 20X/0.8 objective (Zeiss) was used. Multiple images were taken for each animal with multiple z planes of 1 µm interval to get the whole sample thickness. Images were tiled together using Fiji (Schindelin et al. 2012).

Imaging of the whole-mount animals stained for peroxidase activity, actin and nuclei were imaged using an inverted motorized stand Nikon Ti2-E Eclipse equipped with CSU-W1 Confocal Scanner Unit (Yokogawa) with a 50um pinhole disk unit, 2 black-illuminated EMCCD cameras (iXon-Ultra-888 model, Andor) and controlled with VisiView 6.0 imaging software. A Plan Apo lambda oil 40x/1.40 objective (Zeiss) was used.

### Transcription Factor Binding site analysis

Identification of transcription factor binding sites was performed using the JASPAR (Rauluseviciute et al. 2023) collection of transcription binding profiles accessed and analyzed using the JASPAR2018 and TFBSTools Bioconductor packages respectively. Briefly, PWMs of log2 probability ratios were compiled using a pseudocount of 4 and a flat uniform prior as background nucleotide probabilities. The promoter sequence queried encompassed 1500nts upstream of the Transcription Start Site masked for repetitive sequences. Both strands were queried (function TFBSTools::searchSeq) using a minimum score cutoff of 80%. The motifs utilized specifically for the Zic4 and Gata3 binding sites correspond to the JASPAR IDs MA0751.1 and MA0037.3, respectively. *Hydra* promoter sequences were compiled from the NCBI *Hydra* RP 105 assembly. Orthologous promoter sequences in various cnidarians were compiled using the latest available annotations and their corresponding sequence coordinates are provided in Supplementary file 2.

### KDs of Zic4 and Gata3 transcription factors

Gene KDs were performed according to a protocol based on (Vogg et al. 2022). Briefly, 20 to 30 animals per treatment were incubated at 4°C for 1 hour and then transferred into an electroporation cuvette (Gene Pulser/Micro Pulser Electroporation Cuvette 0.4-cm gap, Bio-Rad or electroporation cuvettes 4-mm gap, VWR) and washed twice with chilled Milli-Q water. Residual water was removed, and 200 μl of the small interfering RNA (siRNA) solution was added to each cuvette (siRNAs were ordered as 20–base pair RNA duplexes from Integrated DNA Technologies). Each siRNA was diluted in ddH_2_O to a final concentration of 1.35 μM per treatment. Animals were incubated with the siRNA solution for 5 min. After relaxation of the animals, two pulses (150-V range) were applied for 50 ms (Gene Pulser II with RF module, Bio-Rad). After electroporation, 500 μl of ice-cold recovery medium (80% HM, 20% (v/v) DM) was added to the animals. Animals were carefully transferred into a new Petri dish filled with prechilled recovery medium. The next day, the animals were transferred into fresh HM. Three electroporations were performed in this manner with 1 day for recovery in between successive rounds. To quantify the phenotype, animals were then fed normally (3 times) the following week and were fixed or collected for experiments 7 days (actin/peroxidase staining) or 12 days after the last electroporation (*Nematocilin* staining). For regeneration experiments, animals were bisected at day 2 or day 7 after the last electroporation and left to regenerate for three or six days, before proceeding with foot staining or ISH with Crim-like probe. For qPCR analysis, the animals were fed only twice and collected at day 6 post last electroporation.

### Real-time qPCR verification of RNAi efficiency

KDs of Gata3 and Zic4 were performed as described above. Six days after the last electroporation, samples were collected and lysed in 350 μl of RL buffer with 1% β-mercaptoethanol (Single Cell RNA Purification Kit, Norgen), frozen on dry ice, and stored at −80°C. To establish the KD’s efficiency, RNA was extracted per animal for each condition (n = 8−10). To assess the mutual inhibition, 6 *Hydra* per condition were cut in two segments, representing the oral and aboral parts. The head segment contains the upper one third of the animal and the foot segment contains two thirds of the lower body. Six segments were pooled together before RNA extraction. After collecting all the samples, RNA was isolated as outlined in the instructions for the kit. The concentration and purity of the extracted RNA was verified using a NanoDrop 1000 (Thermo Fisher Scientific). Samples with low RNA quality or insufficient amount of RNA were excluded. Next, cDNA was prepared with the Oligo(dT)12-18 Primer (Thermo Fisher Scientific) using the High-Capacity cDNA Reverse Transcription Kit (Applied Biosystems) as per manufacturer’s instructions. The qPCR reactions were carried out using the StepOnePlus Real-Time PCR cycler (Thermo Fisher Scientific) with a standard run method. Each reaction contained 70 ng (individual animals) or 100 ng (pooled segments) of cDNA, 1× Platinum SYBR Green PCR Master Mix (Thermo Fisher Scientific), and 10 μM primers in a total volume of 25 μl. The sequences of the primer pairs used are given in Table S2. The quantification of gene expression was performed with the Livak method, using GAPDH as housekeeping gene.

### Whole-mount in situ hybridization (*Crim*-like)

In situ hybridization (ISH) was performed according to previously published protocols (Bode et al. 2008) with minor modifications. Briefly, animals were fixed at 4°C in 4% PFA overnight and then dehydrated in a series of 5 min washes in 25-50-75-100% methanol and incubated overnight in methanol at −20°C. Rehydration then followed by successive 10 min washes of 75-50-25-0% methanol in PBST. Samples were then treated for 10 min with a proteinase K solution (10 μg/ml), and the reaction was stopped by incubation with glycine (4 mg/ml) for 10 min. After 2x 5-min washes with PBST and 2x 5-min washes with 0.1 M triethanolamine solution, samples were treated with acetic anhydride for 2x 5 min, washed in PBST (2x 5 min), and refixed in 4% PFA for 20 min. Thorough washing with PBST (5x 5 min) then followed, and the animals were heat treated for 30 min at 80°C. Following preincubation with the hybridization solution for 2 hours at 55°C, digoxigenin (DIG)–labeled RNA probe dissolved in hybridization solution was added. We used the probe for Crim-like as previously published (20) and then washed with a series of 75-50-25-0% solutions of hybridization buffer in 2× saline sodium citrate buffer (SSC). After blocking the samples in 20% sheep serum for 2 hours at 4°C, an incubation with the alkaline phosphatase–conjugated anti-DIG antibody followed (overnight, 4°C). To remove the unbound antibody, 8x 1-hour washes with maleic acid buffer were performed, followed by an overnight wash in the same buffer. The next day, samples were treated by NTMT and levamisole, as detailed in the original protocol. Last, the chromogenic reaction was performed using the bromochloroindolyl phosphate–nitro blue tetrazolium Color Development Substrate (Promega) according to the manufacturer’s instructions.

## Author Contributions

**Jaroslav Ferenc:** Conceptualization, Methodology, Formal analysis, Investigation, Data Curation, Writing – Original Draft, Writing – Review & Editing, Visualization. **Marylène Bonvin:** Methodology, Validation, Formal analysis, Investigation, Data Curation, Writing – Original Draft, Writing – Review & Editing, Visualization. **Panagiotis Papasaikas:** Software, Formal analysis, Data Curation, Visualization. **Jacqueline Ferralli:** Validation, Investigation, Resources, Project administration. **Clara Nuninger:** Validation, Investigation. **Charisios D. Tsiairis:** Conceptualization, Writing – Original Draft, Writing – Review & Editing, Supervision, Project administration, Funding acquisition.

## Acknowledgments

We would like to express our gratitude to Iskra Katic for generating the knockdown animals used in this study. We also thank the FMI Facility for Advanced Imaging and Microscopy for their excellent support and expertise in imaging. Our appreciation extends to all members of the Tsiairis lab for their valuable input throughout the course of this work. Finally, we thank our colleagues Matthew Benton, Helge Grosshans, and Prisca Liberali for their thoughtful comments and constructive feedback on the manuscript. Funding for this work was provided by the Novartis Research Foundation.

