## Supplement for "A transcription factor toggle switch determines differentiated epidermal cell identities in *Hydra*"

**This file includes:**

Figures S1 to S5

Tables S1 and S2

Legends for Supplementary Files 1 and 2

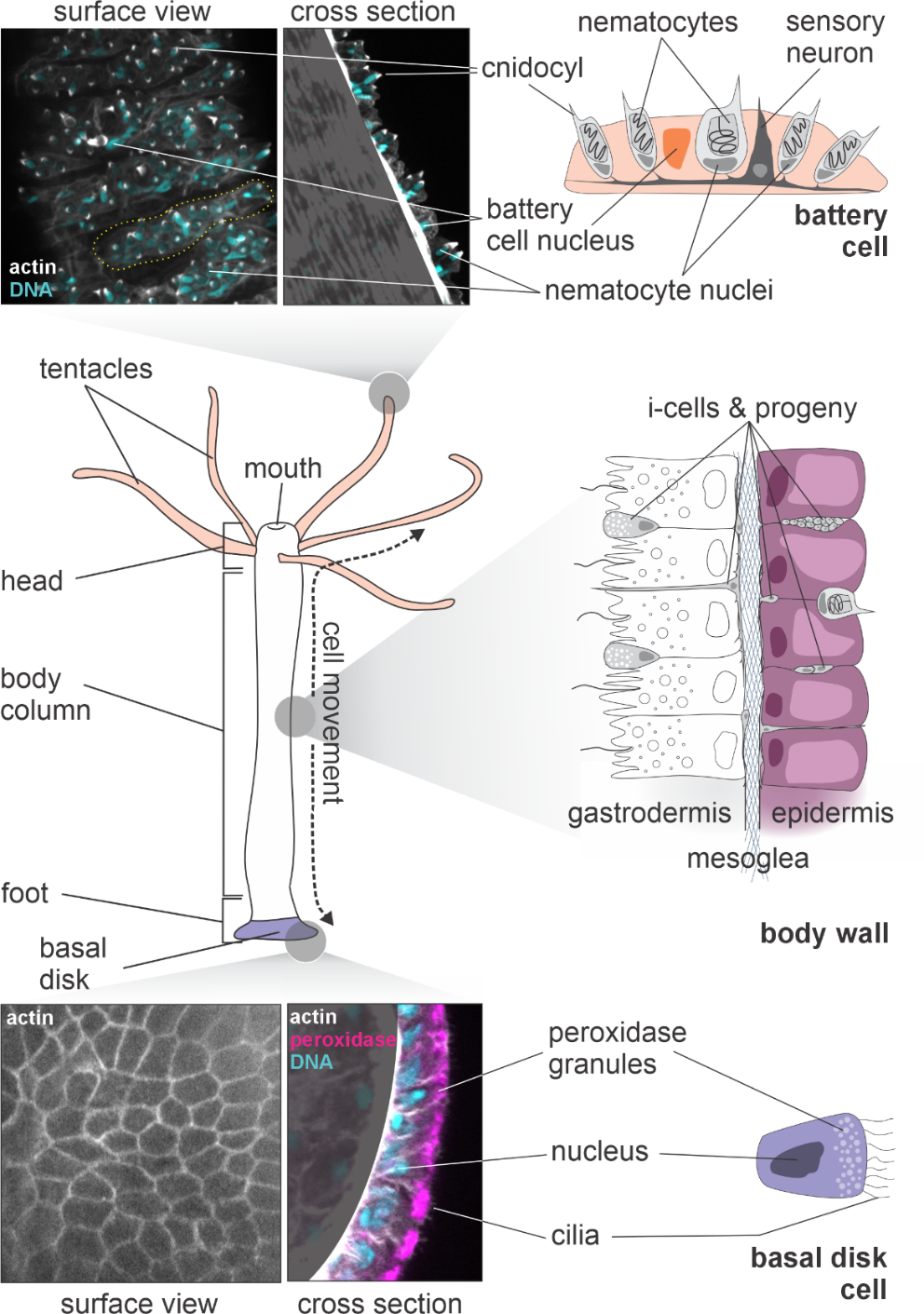

Fig. S1. *Hydra* anatomy and epidermal cell differentiation. The *Hydra* body is organized along a single oral-aboral (head to foot) axis. The body column consists of two layers of epithelio-muscular cells: epidermis and gastrodermis, separated by the mesoglea – a thin layer of extracellular matrix. In between the epithelial cells, i-cells and their progeny (eg. neurons and gland cells) are intercalated. Epithelio-muscular cells of the body wall are not terminally differentiated. They divide and are thereby displaced towards the extremities, where they terminally differentiate. Epidermal cells in tentacles differentiate into battery cells, hosting several nematocytes (stinging cells) in complex with a sensory neuron. Using a DNA staining, a single battery cell (highlighted by a yellow dotted line in the surface view) therefore appears multinucleate. Most of the nuclei however correspond to the associated nematocytes as shown by the proximity of actin bundles characteristic of cnidocils (sensory processes of nematocytes involved in their discharge). At the aboral end, epithelial cells adopt the basal disk identity. Basal disk cells have numerous peroxidase-containing granules close to the apical surface, which are believed to be involved in substrate attachment.

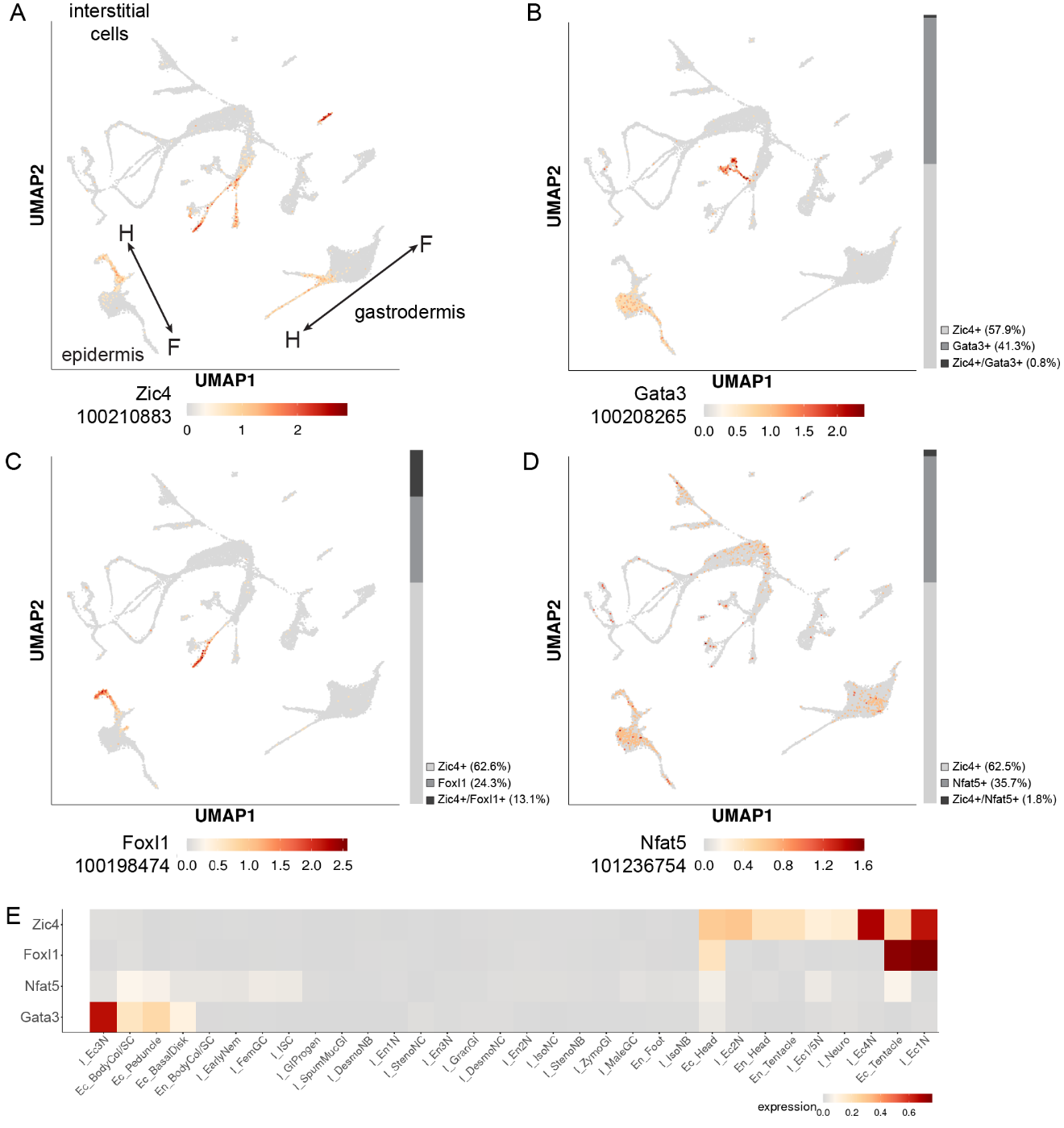

Fig. S2. Expression profiles of candidate TFs in the whole-animal scRNA-seq dataset. (A-D) UMAPs from publicly available scRNA-seq data (research.nhgri.nih.gov/HydraAEP) showing the three Hydra cell lineages (interstitial, epidermal and gastrodermal cells) and the expression profile of candidates TFs. Arrows indicate the direction of the axis between the two poles: oral (head = H) - aboral (foot = F). Grey bar on the right indicates the percentage of cells expressing Zic4, one of the candidate TFSs and double positive cells. (A) Zic4 (100210883/G004456). (B) Gata3 (100208265/G022640) (C) FoxI1(100198474/G021359) (D) Nfat5 (101236754/G023405). For absolute numbers see Table S1. (E) Heatmap showing the expression of the TF candidates across cell types.

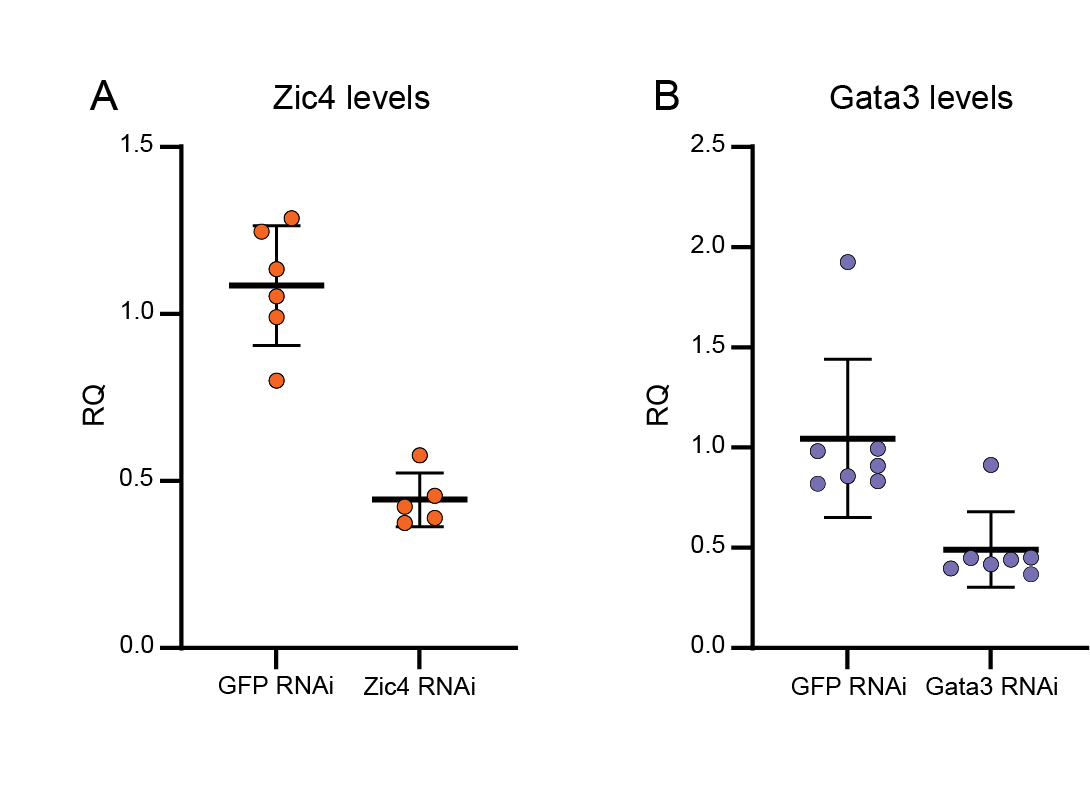

Fig. S3. *Zic4* and *Gata3* KD efficiency. (A) Quantitative PCR analysis of *Zic4* transcript in individual animals after *GFP* RNAi (n=6) and *Zic4* RNAi (n=5). (B) Quantitative PCR analysis of *Gata3* transcript in individual animals after *GFP* RNAi (n=7) and Gata3 *RNAi* (n=7).

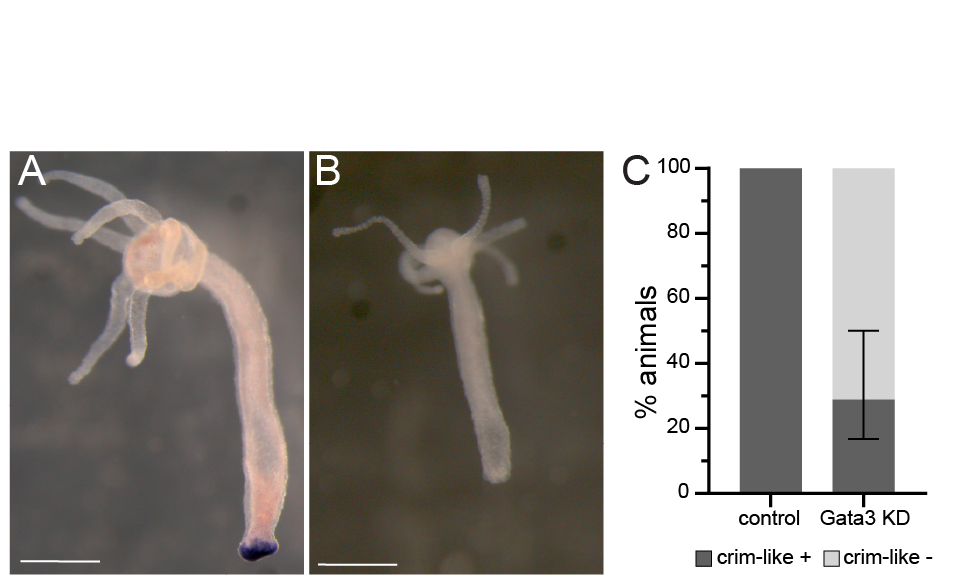

Fig. S4. Expression of the foot-specific marker *Crim*-like in regenerating feet upon *Gata3* KD. (A-B) Whole-mount ISH of crim-like gene, 6 days after bisection in control animals (A) or Gata3 KD (B). (C) Quantification showing the percentage of animals with *Crim*-like expression in the foot at day 6 post bisection, n_control_ = 22, n_Gata3KD_ = 19. Animals pooled from 3 independent replicates. Error bars indicate minimum and maximum values across replicates. Scale bars A-B, 500 μm.

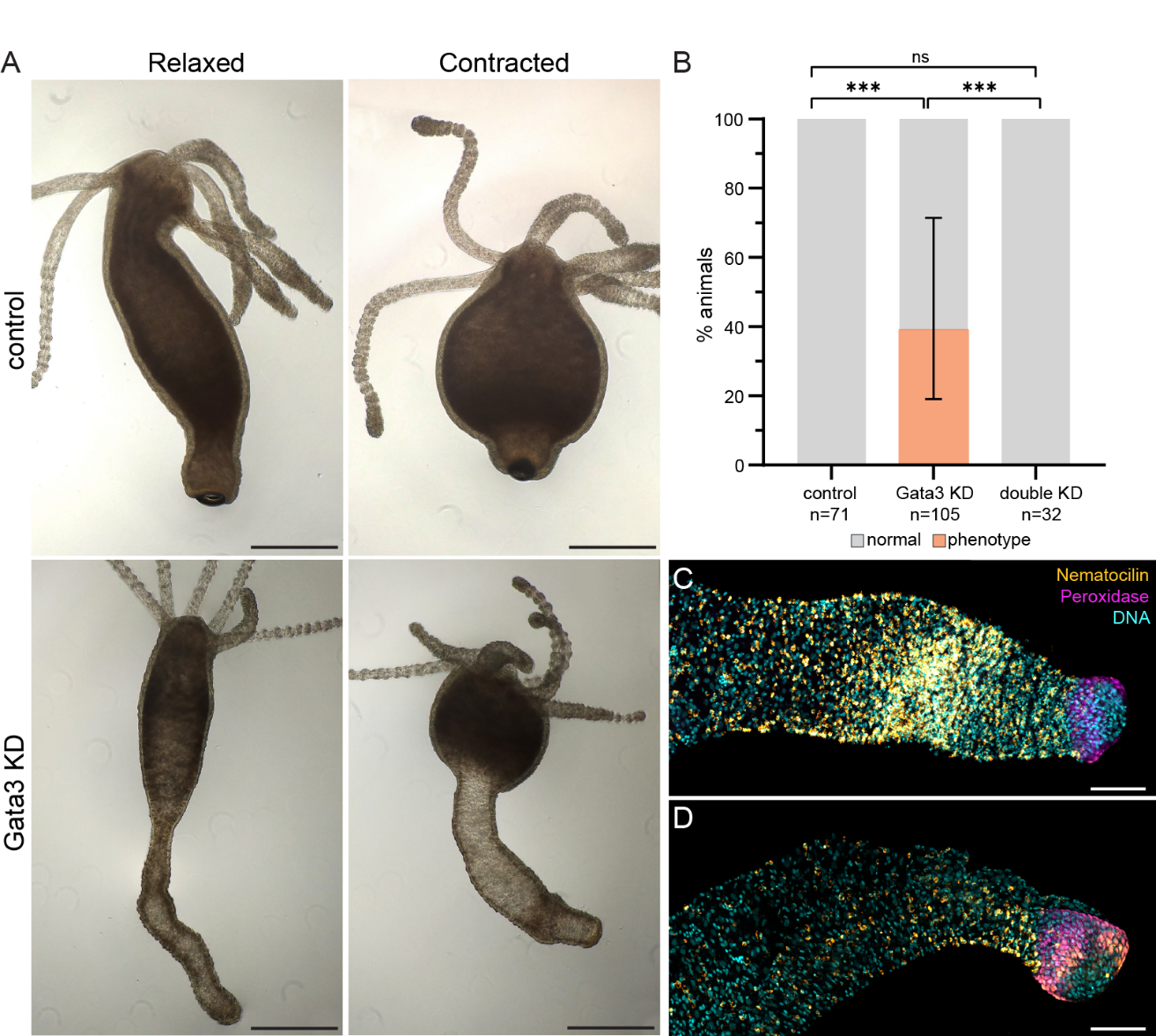

Fig. S5. Varying phenotype strength of the *Gata3* KD. (A) Brightfield images of animals 12 days after last electroporation of *GFP* RNAi (control) or *Gata3* RNAi. *Gata3* KD shows a case of extreme phenotype, with an aboral end transformation into a tentacle-like appendage. Note the inability of this appendage to contract the same way as a foot. (B) Quantification of animals showing the extreme phenotype under different RNAi treatments. Data for n animals pooled from at least 2 independent replicates. Error bars indicate minimum and maximum values across individual replicates. (C-D) Maximum intensity projection confocal images of aboral end upon *Gata3* KD representative of the different phenotypes quantified in Fig.4. (C) Strong phenotype indicates a dense nematocytes patch covering almost the entire area of body wall adjacent to the basal disk (D) Partial phenotype indicates a smaller patch or increased number of scattered nematocytes. *Nematocillin* (orange), Peroxidase (magenta) and DAPI (cyan). Scale bars A-D, 500 μm, F-G: 100 μm.

Table S1. Counts of cells with nonzero expression of candidate TFs. Related to Fig. S2 A-D

|  | ***Gata3*** | | ***FoxI1*** | | ***Nfat5*** | |
| --- | --- | --- | --- | --- | --- | --- |
|  | **nCells** | **%** | **nCells** | **%** | **nCells** | **%** |
| **other TF only** | 884 | 41.26984 | 404 | 24.30806 | 697 | 35.65217 |
| ***Zic4* only** | 1240 | 57.88982 | 1040 | 62.57521 | 1222 | 62.50639 |
| **both** | 18 | 0.840336 | 218 | 13.11673 | 36 | 1.841432 |
| **total** | 2142 | 100 | 1662 | 100 | 1955 | 100 |

Table S2. Q-PCR primers

|  | **forward primer (5' -> 3')** | **reverse primer (5' -> 3')** |
| --- | --- | --- |
| ***Gapdh*** | AAATGGCAAGCTAACTGGAATGG | TCAGATGCAGCTTTTACTTTGGC |
| ***Gata3*** | TAAACCAAAGAGGAGATTGTCAC | ATACTGGTTCACCACTTCCA |
| ***Zic4*** | GGAAAATACCCAAAGCACGA | TATCCGTACTGTGGGCTTCC |

**Supplementary File 1 (separate file).** **Transcription factor enrichment analysis.**

*TF_enrichment_Fspecific:* summarizes the results of enrichment analysis in the foot-specific genes upregulated in tentacles upon *Zic4* KD. For each factor, JASPAR ID, name, class, fraction of genes with binding sites in the dataset, fold enrichment over all expressed genes, and adjusted P value are given.

*Zic4onTFpromoters:* all detected *bona fide* Zic4 binding sites in the transcription factor promoters across the *Hydra* genome. For each gene, NCBI GeneID is given along with the blast hits for closest vertebrate ortholog and corresponding annotation. The position of Zic4 binding sites is indicated relative to the TSS.

**Supplementary File 2 (separate file).** **Zic4 and Gata3 binding sites conservation analysis.**

*Sequences:* NCBI sequence coordinates for all sequences used in the analysis. These sequences correspond to the segment 1.5 kb upstream of the TSS. If a given gene has several transcript variants with different TSS, the sequence begins immediately upstream of the proximal-most TSS, extending to 1.5 kb upstream of the distalmost TSS (relative to the coding sequence).

*TSScoordinates:* Coordinates of all unique TSS in the sequences

*GATAonZIC:* All *bona fide* Gata binding sites in *Zic* gene promoters. For each binding site, the species and transcript isoform are indicated along with the site sequence and strength, as well as coordinates relative to the respective TSS.

*ZIConGATA:* All *bona fide* Zic binding sites in *Gata* gene promoters. Same information given as for the Gata sites.
